# Progress towards lymphatic filariasis elimination in Ghana from 2000-2016: analysis of microfilaria prevalence data from 430 communities

**DOI:** 10.1101/507145

**Authors:** Nana Kwadwo Biritwum, Kwadwo K. Frempong, Suzanne Verver, Samuel Odoom, Bright Alomatu, Odame Asiedu, Periklis Kontoroupis, Abednego Yeboah, Edward Tei Hervie, Daniel A. Boakye, Sake J. de Vlas, John O. Gyapong, Wilma A. Stolk

## Abstract

**Background:** Ghana started its national programme to eliminate lymphatic filariasis (LF) in 2000, with mass drug administration (MDA) with ivermectin and albendazole as main strategy. We review the progress towards elimination that was made by 2016 for all endemic districts of Ghana and analyze mf prevalence from sentinel and spot-check sites in endemic districts.

*Methods:* We reviewed district level data on the history of MDA and outcomes of transmission assessment surveys (TAS). We further collated and analyzed microfilaria (mf) prevalence data from sentinel and spot-check sites.

*Results:* MDA was initiated in 2001-2006 in all 98 endemic districts; by the end of 2016, 81 had stopped MDA after passing TAS and after an average of 11 rounds of treatment (range 8 – 14 rounds). The median reported coverage for the communities was 77-80%. Mf prevalence survey data were available for 430 communities from 78/98 endemic districts. Baseline mf prevalence data were available for 53 communities, with an average mf prevalence of 8.7% (0 - 45.7%). Repeated measurements were available for 78 communities, showing a steep decrease in mean mf prevalence in the first few years of MDA, followed by a gradual further decline. In the 2013 and 2014 surveys, 7 and 10 communities respectively were identified with mf prevalence still above 1% (maximum 5.6%). Two stopped MDA in 2015 and 2016 respectively, while the rest of the 15 communities above threshold are all within 13/17 districts where MDA is still ongoing.

*Conclusions:* The MDA programme of the Ghana Health Services has reduced mf prevalence in sentinel sites below the 1% threshold in 81/98 endemic districts in Ghana, yet 15 communities within 13 districts (MDA ongoing) had higher prevalence than this threshold during the surveys in 2013 and 2014. These districts may need to intensify interventions to achieve the WHO 2020 target.

*Author summary:* Lymphatic filariasis (LF) control in Ghana has relied on ivermectin and albendazole since the year 2000 when the Ghana Filariasis Elimination Programme started. We analyzed trends in microfilaraemia prevalence during MDA, reported coverage, and transmission assessment survey using data obtained from the Ghana Health Services (GHS). The median reported treatment coverage varied between 77-80% over the years. Our results show that the treatment in Ghana made a significant impact in reducing infections <1% in majority of sentinel sites in endemic districts (81/98) by 2016. In the remaining 17 districts, extra efforts may be needed to achieve the same goal. Some of the challenges could be low coverage in some communities, high baseline endemicity, programme logistical challenges etc. The required average rounds of MDA needed for elimination was 11, higher than that proposed by the Global Filariasis Elimination Programme. This article is relevant to LF control programmes in assessing the impact of MDA. It is important for programmes to monitor infections especially within communities where mf prevalence is still above the 1% threshold to ensure that the WHO 2020 elimination target is achieved.

## Introduction

Lymphatic filariasis (LF), commonly known as elephantiasis, is a debilitating and disfiguring tropical disease caused by lymphatic-dwelling filarial parasites *Wuchereria bancrofti, Brugia malayi* and *Brugia timori.* The disease is transmitted by different species of mosquitoes depending on the geographical location, including *Culex, Anopheles* and *Aedes* species. About 90% of the worldwide cases are caused by *W. bancrofti* and 10% caused by *B. malayi* and *B. timori.* Based on re-assessment of the global prevalence and distribution of LF [1], more than 120 million people were found to be infected and 40 million disfigured and incapacitated in the year 2000 [2]. In the same year, the Global Programme to Eliminate Lymphatic Filariasis (GPELF) was established, aiming to eliminate the disease as a public health problem by 2020 through annual mass drug administration (MDA) with albendazole in combination with diethylcarbamazine citrate (DEC) or ivermectin to all individuals at risk [3].

By the end of 2016, 20 out of 73 countries originally listed by WHO as being endemic for LF have stopped interventions after passing the first transmission assessment survey and are conducting surveillance to validate elimination. Additional 30 countries have delivered MDA at least once in all endemic areas and are also on track to achieve elimination [4]. While many have passed the TAS, there are also reports of failure [5] and of ongoing transmission in spite of passing the TAS [5-7].

A national survey carried out in Ghana in 1994 showed that the microfilaraemia prevalence varied from 0 - 20% between regions [8]. In the highly-endemic Kassena Nankana district (Upper East Region of Ghana), the prevalence of hydrocele was 30.8% and elephantiasis of the leg was 3.8% in the population aged 10 years and above [9,10]; 12% of extended families reported to have at least one family member with elephantiasis of the leg [10]. The extensive mapping of endemic communities [11] provided a database on areas in Ghana and neighboring countries that needed more efforts to eliminate the disease.

The LF elimination programme in Ghana started in 2000 and gradually scaled up over the years and by 2006 all endemic districts were covered. The implementation and outcomes by district were described in two recent papers [12,13]. By 2016, 81 of 98 initially endemic districts had reached an mf prevalence <1%, had passed TAS survey and stopped MDA, while the remaining districts still had mf prevalence >1% [13] in spite of at least 10 years of MDA. The required duration of MDA turned out to be longer than the anticipated 5-6 years, which might be due to relatively high baseline mf prevalence levels. There were no major differences with other districts in reported coverage of MDA or long-lasting insecticide treated bednets [13].

Expected trends in infection during MDA will depend on multiple factors, including local baseline endemicity (depending on local transmission conditions) and the achieved coverage and compliance with MDA. To obtain better understanding of these factors, in this paper we present and analyze community-level data from microfilaraemia prevalence surveys and transmission assessment surveys (TAS) from sentinel and spot-check sites for all endemic districts in Ghana. So far, this study represents the longest and largest LF programmatic study in Africa.

## Methods

### Ghana Filariasis Elimination Programme

The Ghana Filariasis Elimination Programme (GFEP) was established in June 2000 following the establishment of the Global Programme to Eliminate Lymphatic Filaraisis. Mapping of communities started in 2000 using the 50-km sample grid, rapid assessment procedure for antigenaemia in sample villages and spatial analysis to plot prevalence contours from 2000 to 2001 [11,14]. Forty nine districts were initially identified as endemic and therefore selected for implementation of MDA. The GFEP implementation, programme outcomes, challenges and districts re-demarcation have been described in Biritwum *et al.* (2017a). Based on current demarcation, 98/216 districts (45%) are endemic with LF in Ghana.

The treatment implemented in Ghana was the combination of ivermectin (150 μg/kg) and albendazole (400 mg) given annually by the community-directed treatment approach [15] and implemented at the district level. MDA usually took place between March and June in all endemic communities across the country. Individuals eligible for treatment were those aged ≥5 years (excluding pregnant women, lactating mothers and the sick), and selection was solely based on height (≥90 cm) for those whose ages were not known. MDAs usually lasted for about 1 - 2 weeks per community. Individual treatment information (whether treated, absent, pregnant, sick, etc) was recorded in the community treatment book and summarized into treatment records by the Ghana Health Services (GHS). Community-level treatment coverage data (number treated out of total population at risk) across the country were reviewed and summarized by the GHS. For the purpose of this study, summary reports were reviewed. There was no treatment offered in 2011 due to logistic and funding challenges; 2009 and 2012 treatment data were not available.

### Monitoring and evaluation

#### Parasitological surveys

Parasitology data were collected by the GHS in programmatic yearly surveys (2000 - 2014) in selected endemic communities. In 2000, baseline mf surveys were carried out in 24 purposefully selected endemic communities (based on known high endemicity and population stability) [16,17] from the 8 districts where MDA was first initiated. From 2001-2004, baseline mf surveys were done in sentinel sites of remaining districts, as MDA was being extended into these districts. Subsequently, the previously selected sentinel sites per district were repeatedly surveyed to monitor progress to elimination (usually once every 2 or 3 years, but sometimes the interval was much longer due to financial constrains). Additional surveys were done in spot-check sites (same characteristics as sentinel) that were surveyed only once and often selected randomly from the same district where sentinel site is located to cross check the MDA performance in that district.

The mf surveys were usually done at the end of the year between November and December (before MDA treatment was done the following year between March-July). The target number of persons for sampling increased over time based on WHO guidelines; between 2000 - 2002 the target was 100 persons per community, between 2003 - 2009 it was 500 persons, and after 2009 it was 1000 - 1500 persons. The surveys were usually preceded by a community gathering or announcement by the team informing members of the community to converge for the night blood collection (9pm - 2am). Those selected for sampling were verbally consenting individuals (parent consented for their children) aged ≥ 5years or with height ≥ 90cm height for those whose ages were not known, including pregnant women and lactating mothers. Blood was sampled from these individuals by finger pricking (middle or forth finger) and a volume of 60μl taken for thick blood smear test (2000 - 2009) [18]. In later years (2010 - 2014), the volume of blood sampled was increased to 100μl and the microfilariae were counted using a regular microscope with a rafter counting chamber [19], by trained GHS laboratory technicians.

#### Transmission assessment surveys

The transmission assessment surveys (TAS) in Ghana followed the WHO guidelines, using antigenaemia prevalence in children aged 6 - 7 years as indicator of active transmission of LF [20]. TAS has a different sampling system and a different target population than a full mf survey. In Ghana, the elimination programme used a district as an implementation unit (IU) for MDA. The evaluation unit (EU) to assess progress of programme may also be a district or a cluster of districts with a population not more than 2 million. In some of the cities where the district population was more than 2 million, the district was divided into different EUs. An EU qualified to be assessed after achieving treatment coverage of ≥65% for 5 years and also recording mf prevalence of <1%. Those EUs who met the criteria were selected for the TAS. The TAS involved sampling of children 6 - 7 years in primary schools within the EU after written consent from their parents. The schools to be surveyed, the number of children to be tested and the critical cut-off point (maximum number of positives to fail TAS) were estimated using a survey sample builder software recommended by WHO [21]. A volume of 100 μl blood was taken from the children by finger pricking and test done using immunochromatographic card test (ICT) [22]. The EUs where the number of positive children was less than the critical cut-off point, passed the TAS (TAS-1) and the MDA was stopped after the last treatment. The MDA stop decision is based on TAS-1. After stopping MDA, two more TAS surveys (TAS-2 & TAS-3) are done after 2 - 3 years and 5 years respectively before elimination said to be achieved. During TAS-1, the EU with number of positive children above the critical cut-off point, failed the TAS and continued MDA for 2 - 3 years. In such EUs, a community survey was required to achieve an mf prevalence of <1% before TAS-1 was repeated [13].

### Data collation and analysis

The longitudinal parasitological and treatment data from 2000 - 2014 were collated along with background information from the GHS and updates on TAS results till early 2016. Parasitological data comprised of number examined and number that were microfilarial positive in each community. Community mf prevalence was estimated as the number of microfilarial positives as a percentage of number examined. Mean mf prevalence (for districts and country) were calculated by estimating the percentage of total positive / total examined and total treated / total population of communities included in the sample respectively. We present data on community, district and country level.

#### Data limitations and special situations

We only consider mf prevalence data in our analysis of community-level trends in infection prevalence. No mf prevalence data were available for 2001 and data for 2002 were limited to few sites, as all community surveys in 2001 and part of the survey in 2002 only used ICT antigenaemia tests. For 3 districts in 2003 (with 2 communities surveyed per district) and all districts in 2005 (with 3-4 communities surveyed per district) mf prevalence was not reported for each community separately, and we only know the overall mf prevalence - aggregated over the surveyed communities. These aggregated data points were included in combined trends (averages) for all communities, but were not matched to community specific data for analysis for time trend analysis. There were no mf surveys carried out in 2006. Mf prevalence data in 2008 and 2010 were excluded from trends and analysis since communities sampled in these years were not randomly selected (mf data from individuals closely related to school children who were positive using ICT during school surveys).

### Ethics Statement

Ethical clearance was obtained from the Ghana Health Service Ethical Review Committee (ID NO: GHS-ERC-10/0/06) and the Liverpool School of Tropical Medicine’s Research Ethics Committee’s Research Protocol Approval (06.47). The study obtained oral informed consent from adult participants while parents and guardians orally consented for their children and wards to be part of this study. Due to the programmatic nature of the study with regular MDA and mf surveys done in many sites, participants in these communities were aware of the program. Given that the communities were mainly rural with study participants having minimal or no education and being suspicious of signing documents they did not well understand, oral consent was applied and noted as part of questionnaires during the surveys. Oral informed consent was approved by the Ghana Health Service and Liverpool School of Tropical Medicine’s ethical review committees.

## Results

### Overview of MDA implementation in Ghana

MDA started between 2000 and 2001 in 10/98 districts selected from the northern and coastal regions of Ghana. In 2002, 17 more districts were enrolled onto the MDA programme and in 2003, 2004 and 2005 a number of 27, 16 and 25 more districts were enrolled onto the MDA programme respectively. By 2006 all the 98 endemic districts had been enrolled (Figures 1 and 2, Table 1). All communities in each district were expected to be treated in the same year MDA started, thus geographical coverage within a district was expected to be 100%.

**Table 1:** Assessment of mf prevalence.

| **Assessment of mf prevalence** | **2000** | **2001 \*** | **2002** | **2003 \*** | **2004** | **2005 \*** | **2006 \*** | **2007** | **2008 \*** | **2009 \*** | **2010 \*** | **2011** | **2012 \*** | **2013** | **2014** | **2015** | **2016** |
| --- | --- | --- | --- | --- | --- | --- | --- | --- | --- | --- | --- | --- | --- | --- | --- | --- | --- |
| No. of IUs that started MDA | 1 | 9 | 17 | 27 | 16 | 25 | 3 | 0 | 0 | 0 | 0 | 0 | 0 | 0 | 0 | 0 | 0 |
| No. of IUs that stopped MDA | 0 | 0 | 0 | 0 | 0 | 0 | 0 | 0 | 0 | 0 | 5 | 0 | 0 | 0 | 65 | 9 | 2 |
| No. of communities included in mf prevalence surveys (No. of districts) | 24 (8) | 0 (0) | 10 (5) | 40 (18) | 25 (12) | 34 (10) * | 0 (0) | 59 (9) | 0 (0) | 111 (21) | 0 (0) | 109 (31) | 103 (19) | 24 (13) | 74 (16) | - | - |
| No. of people examined for mf | 2,607 | – | 1,784 | 4,603 | 2,933 | 4,579 | – | 7,643 | – | 15,175 | – | 15,675 | 19,268 | 11,026 | 13,901 | - | - |
| Average mf prevalence (%) in examined communities (range) | 23 (11-46) | – | 2.8 (0-12) | 7.1 (0-28) | 8.1 (0-34) | 2.0 (0-4) | – | 3.5 (0-21) | – | 2.4 (0-18.8) | – | 0.23 (0-15) | 0.68 (0-9.8) | 0.85 (0-3.7) | 0.57 (0-5.6) | - | - |
| No. of communities with mf prev >1% (%) | 24 (100) | – | 5 (50) | 25 (62.5) | 17 (68) | 6 (60) | – | 37 (62.7) | – | 44 (39.6) | – | 3 (2.8) | 18 (17.5) | 6 (25) | 10 (13.5) | - | - |

IU = implementation Unit, MDA = mass drug administration, Com=community, Cov=coverage, prev=prevalence, mf=microfilaraemia, No. = number
\*Refer to Data limitations and Special situations above.

**Fig 1:**
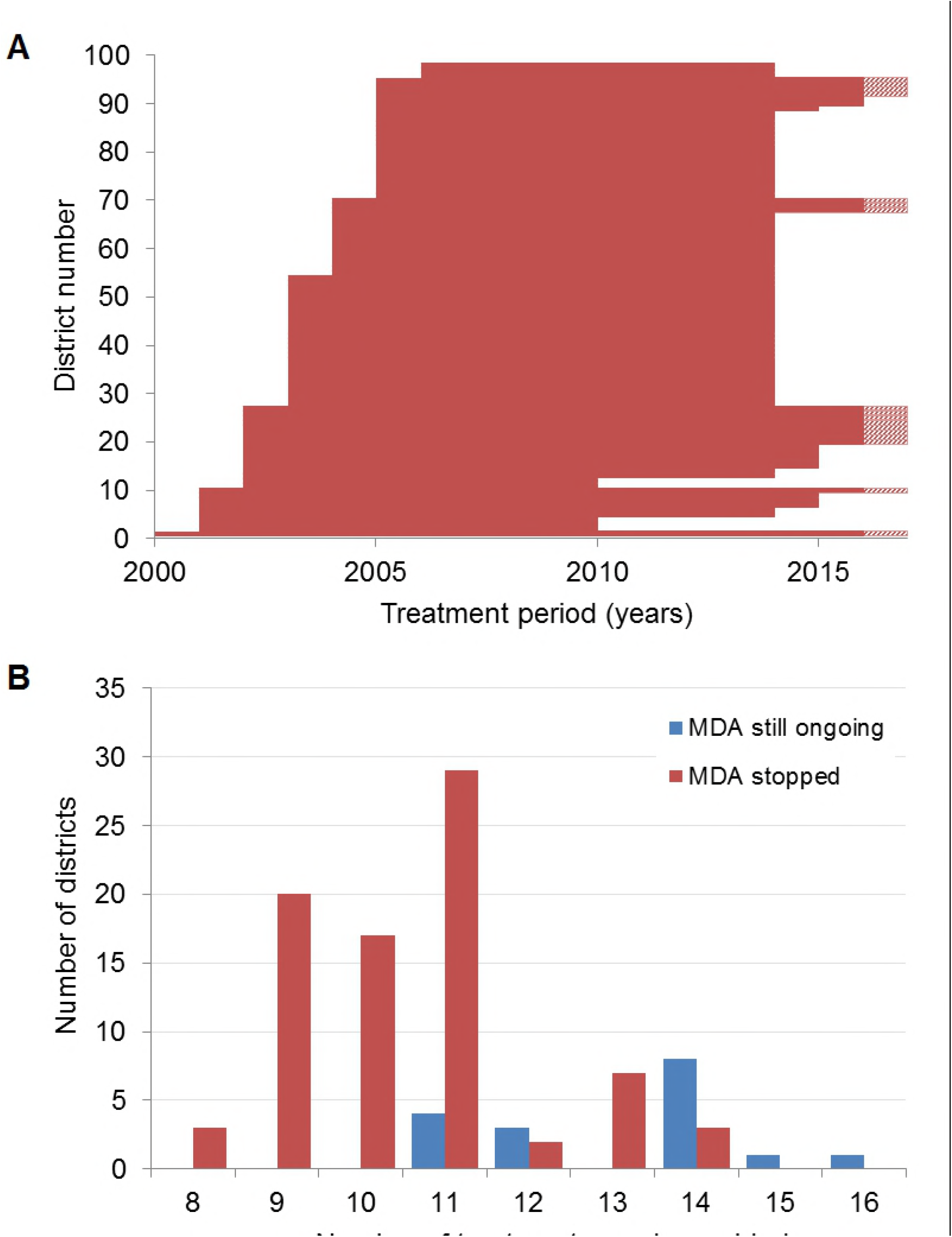
Implementation of district-level MDA in Ghana. A) Period of MDA by each district in order of start year. Each horizontal line represents a district. Bars with a dashed section on the right-hand side represent districts where MDA is still ongoing after 2016 with unknown end year. See supplementary Table S2 for more details. B) Frequency distribution of the number of treatments provided by district through 2016, presented separately for districts that had stopped MDA by 2016 and those with still ongoing MDA.

**Fig 2:**
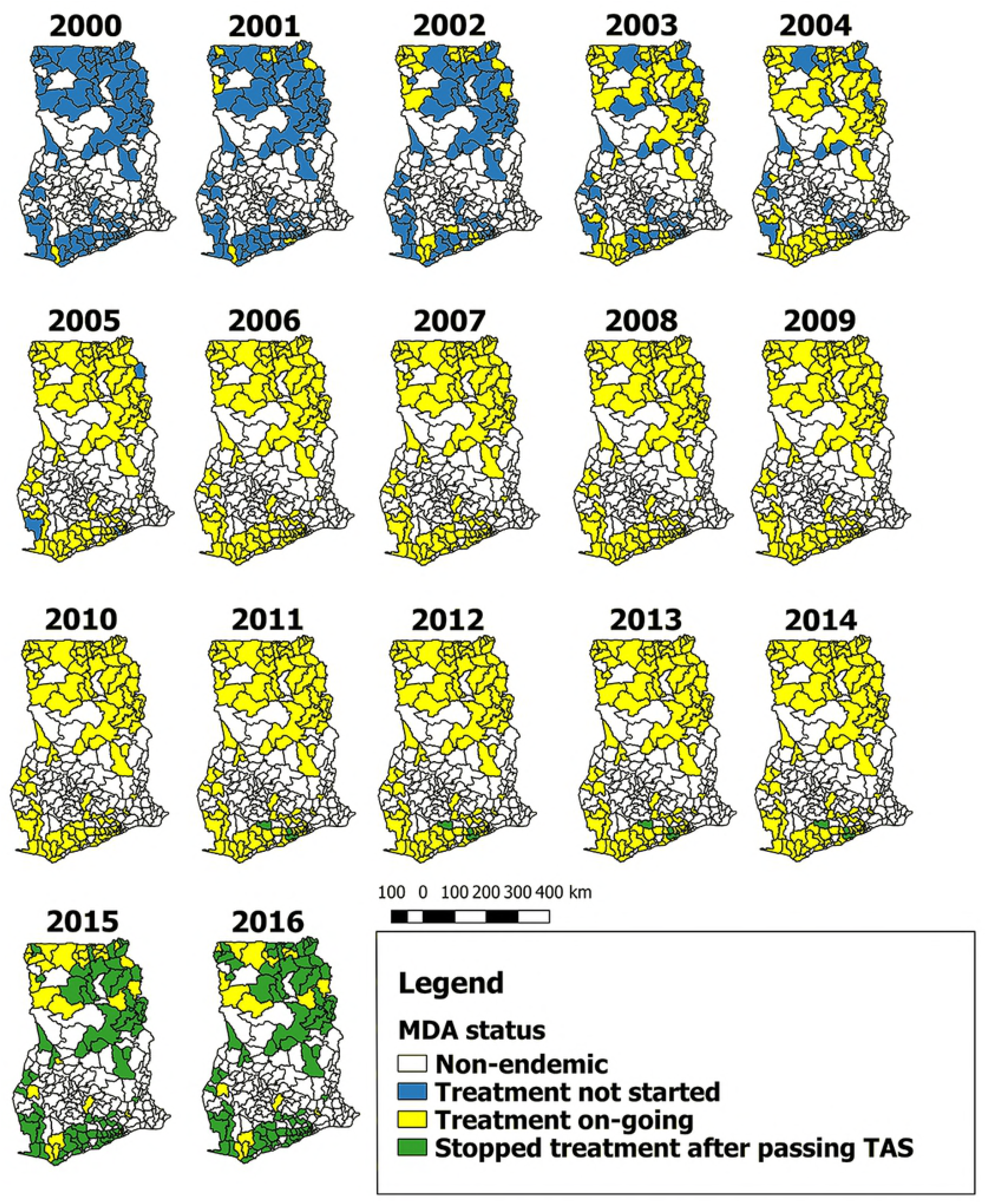
Progress of MDA implementation in Ghana. NB: In the year 2011, there was no treatment due to some logistical challenges. The maps give an overview of the treatment progression to cover all the endemic districts in Ghana.

#### Reported coverage of MDA by calendar year in Ghana

The median reported treatment coverage in treated districts of Ghana seemed to be constant over time, around 77 - 80% between 2000 – 2010, and the interquartile range and distribution of outliers are also similar over time (Figure S2). Although mean reported coverage per year seem to be high, there are large differences between communities. Community-level coverage estimates varied from 10 to 120%, with at least 7952/41265 (19.3%) surveys having a coverage under 65% and 198/41265 (0.5%) surveys over 100%, indicating wrong denominators.

TAS was done in 5 districts in 2010, and all passed. Another 65, 9 and 2 districts had their first TAS in 2014, 2015 and early 2016, respectively and all passed. By the end of 2016, 81 out of the 98 endemic districts had passed the TAS in Ghana and had stopped MDA (Figures 1 and 2). 17 are left, of whom 4, 3, 8, 1 and 1 district have done 11, 12, 14, 15 and 16 rounds of MDA, respectively (see Table S2, supplementary data for details). The average number of treatment rounds in districts that stopped MDA was 11 rounds, varying from 8 – 14. In 6 communities that were re-surveyed in 2014 for mf prevalence after stopping MDA in 2010, mf prevalence was always below <1%. TAS-2 was performed in 69 districts in 2012 or 2015, and all districts passed. Details of the TAS surveillance in Ghana are given in Table S2, supplementary data.

### Trends in Mf prevalence

Mf prevalence data were available from 613 community mf surveys (datapoints), carried out between 2000 to 2014 in 430 communities (292 sentinel sites; 138 spot-check sites) in 78 out of the 98 endemic districts (within 8/10 regions of Ghana). Twenty districts were not represented in our compiled database, either because only antigenaemia data or TAS data were available or because no surveys had been done after re-demarcation of districts. 352 communities were measured only once and 78 measured multiple times (sampled between 2-6 times). Out of those measured multiple times, 35 communities also had data including baseline. Most of the single time point surveys were observed after 2007 (Figure 3A). Overall, the total number of individuals sampled per year ranged between 1,784 – 19,268 (Table 1).

**Fig 3:**
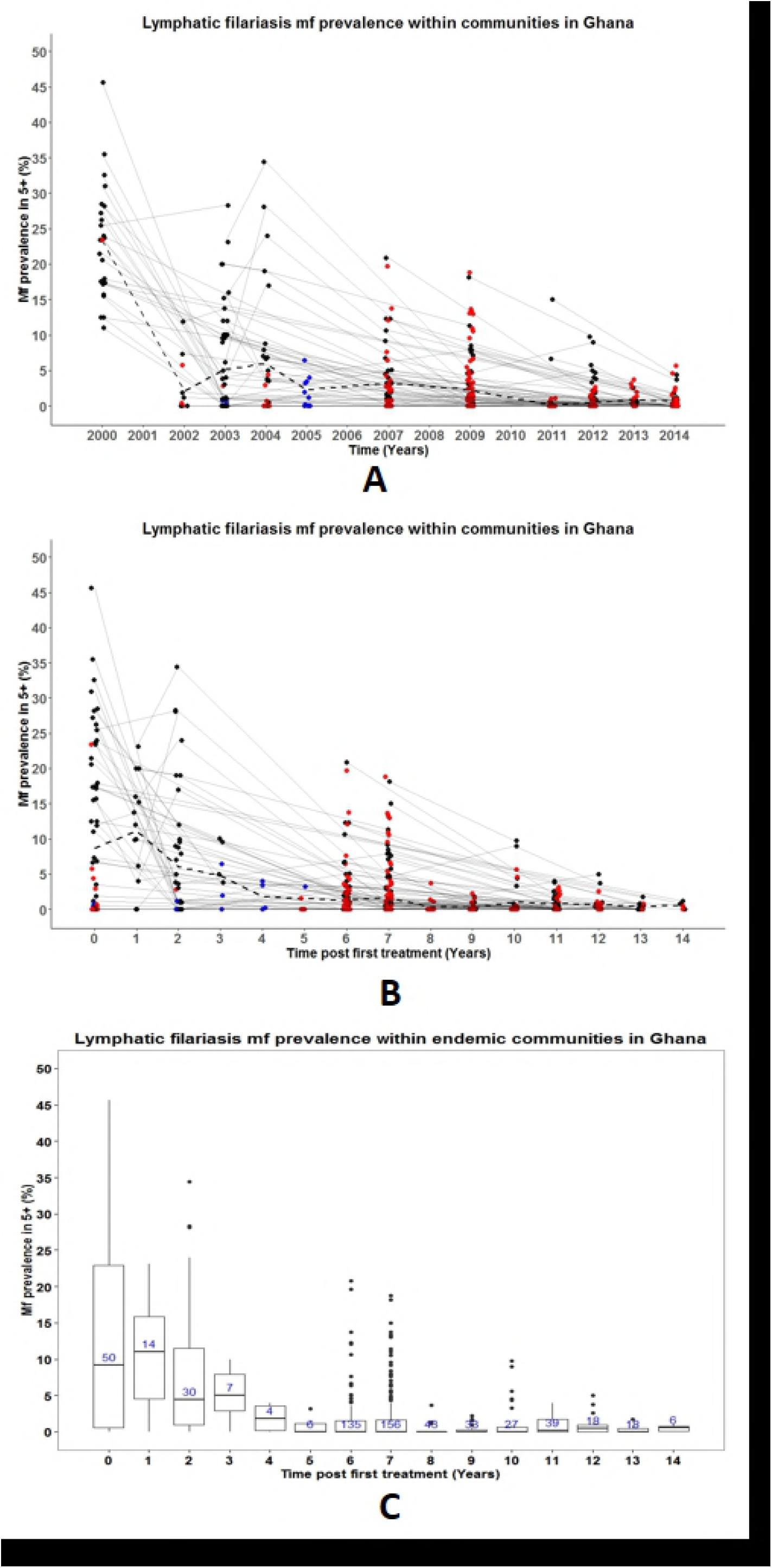
Observed lymphatic filariasis mf prevalence in sentinel and spot-check sites in Ghana, measured in the population aged 5 and above, for the period 2000-2014. A) Data presented by calendar year. Multiple observations from the same community are connected through thin grey lines. Observations from communities surveyed only once are highlighted in red. Observations presenting aggregated prevalence over multiple communities are displayed in blue (in 2003 and 2005). Dashed lines represent the average prevalence from all surveyed communities at each time point. Bullets at the same time point have been jittered to avoid overlapping of points at the same position; these do not represent time in months. B) As panel A but with time since first treatment on the horizontal axis. C) As B, but with data summarized in boxplots. The box at each time post treatment represents the interquartile range of mf prevalence in ≥5 years and the thick horizontal lines across each box represent the median mf prevalence. The bullets outside each box (above or below) represent the outliers and are defined as 1.5 times the interquartile range above or below the ends of the box (25^th^ and 75^th^ percentile). The vertical lines (whiskers) extend to the first value (mf prevalence) before the outlier cut-off and where there are no outliers, they represent the minimum and maximum mf prevalence at each time post treatment. The numbers in the boxes are the total number of communities examined at each time post treatment.

#### Baseline mf prevalence

Baseline parasitological surveys were carried out in the years 2000 - 2004, before the start of MDA, examining 7,882 individuals from 53/430 communities within 21/98 districts. The number of individuals examined per community at baseline ranged between 52 - 441 (mean 137, median 112). The average mf prevalence at baseline was 8.7% (range 0-45.7%) with the highest recorded in Gyahadze located in the central region of Ghana (See supplementary Table S1).

#### Trends in mf prevalence over time (2000-2014)

Community-level mf prevalence data are presented by calendar year (Figure 3A). The impact of MDA on mf prevalence cannot clearly be seen from this figure, due to the differences between communities in start year of MDA. In Figure 3B, therefore, the same data are presented by time since first treatment, while Figure 3C presents these data in boxplots to better visualize the distribution of the observed community-level mf prevalence. From these data, we conclude that the variation in baseline prevalence was huge. The mean and median mf prevalence in surveyed communities declined strongly with increasing duration of MDA. The small increase in median prevalence observed 1 year after the onset of sampling is a selection effect and does not indicate a lack of impact, because surveys were only done in districts with a relatively high baseline mf prevalence. Although 6-7 years after the onset of MDA the median prevalence had fallen below 1%, there was still huge variation between communities and many communities still had mf prevalence levels above 5%. The number of districts and communities surveyed declines over time, because districts that have stopped MDA are no longer included in surveys. In addition, surveys were selectively performed in communities in districts with low reported coverage or relatively high prevalence in previous surveys. For these reasons, trends in mean and median mf prevalence of surveyed villages during later stages of the control are difficult to interpret. Yet, we still see a continued decline in the maximum observed prevalence levels with increasing duration of MDA. In most communities with multiple measurement the mf prevalence steadily decreased over time, but 12 out of 78 (15%) communities had at least once an increase between 2 time points (Figure 3A & B). In the 2013 and 2014 surveys, 7 and 10 communities respectively were identified with mf prevalence still above 1% (maximum 5.6%). Only 2 stopped MDA in 2015 and 2016 respectively. The rest of the 15 communities above threshold are all within 13 out of the 17 districts where MDA is still ongoing.

In 34 districts, one or more communities were surveyed at least twice during the period of MDA. Data for these districts are shown in supplementary file Figure S1. When community data were aggregated at district level, there was a general decrease in average mf prevalence over time to approach zero in most districts (Figure S1, red line). In 4 districts (Bongo, Jirapa, Lambussie-K and Lawra) there were slight increases in mf prevalence after baseline before decreasing steadily. Almost all the districts we assessed, apart from two (Lawra and Wa-West), showed mf prevalence less than 5% after 6 years of MDA (Figure S1, supplementary data). In 31 out of these 34 districts (91%), mf prevalence eventually fell below <1% after 6 - 14 rounds of treatment; this was not the case in three districts (Bole, Jirapa and Wa-West) where the mf prevalence was still ≥1% in 2013 or 2014 and MDA is still ongoing. 51 out of the 78 examined districts/IUs (65%) needed more than 6 rounds of MDA to reach mf prevalence of <1%.

## Discussion

Ghana has made good progress towards elimination since the start of its elimination programme in 2000. The baseline mf prevalence in sentinel sites was 8.7% on average, ranging from 0 to 45.7%. The mf prevalence declined steeply during the first few years after starting MDA in communities, followed by a more gradual decline thereafter (Figure 3A). Surveys performed after 6-7 rounds of MDA showed high variation between communities in mf prevalence, with the mf prevalence often exceeding 1% or even 5% (Figure 3B & C). By the end of 2016, 81 out of 98 endemic districts had stopped mass drug administration (MDA) after an average of 11 rounds of treatment (range 8 – 14). Currently, treatment is still ongoing in 17 districts in Ghana.

We have created a unique longitudinal database on the long-term impact of MDA for lymphatic filariasis (LF) elimination in Ghana, containing data from 430 sentinel and spot-check sites. There are at least 12 countries that have reported longitudinal trend data on at least 3 microfilaria (mf) prevalence surveys of LF after at least 3 rounds of MDA, of whom 5 in Africa: Tanzania, Kenya, Nigeria, Egypt and Mali [23-28]. These African studies have reported the impact of 4 - 10 rounds of MDA on antigenaemia/mf prevalence within 4 - 20 sentinel/study sites where about 50 - 2000 participants were tested per year. Since we have more data (15 years MDA, 430 communities, 1,784 - 19,268 participants), this gives us more insight into the impact of MDA on mf prevalence, the dynamics involved over a period of time and the variability in outcomes between sites.

At country-level we observed huge variation in baseline endemicity level and trends towards elimination (Figure 3 A&B). Patterns became clearer with less variation within districts when we plotted and analyzed data by districts (Figure S1). For some districts only few observations were available, especially in districts with relatively low baseline prevalence, where elimination was relatively rapidly achieved, obviating the need for further surveys. In districts with relatively high baseline mf prevalence, sometimes many rounds of MDA were needed to ensure that mf prevalence reach below 1%.

When GPELF was initiated, it was expected that elimination could be reached after 5 or 6 rounds of MDA with good coverage – ≥65% [3,29]. Although few countries were indeed able to reach elimination within the 5 - 6 years of MDA [2], the required treatment duration in Ghana was always longer, often considerably longer. This experience can help other African countries with planning their interventions. Previous modelling studies already suggested that 5-6 rounds of MDA would not be enough in case of low coverage and/or high baseline endemicity [7,30-32], and the same factors may explain why the required treatment duration in some Ghanaian districts is much longer than in others [13].

Our data confirm the importance of baseline endemicity for the required treatment duration, but the role of coverage was more difficult to proof. Reported coverage data at community level were collated for all endemic district of Ghana (see supplementary data; Figure S2 for details). Although the reported coverage was good for the majority of communities, coverage levels <50% are also frequently encountered. However, such data are notoriously unreliable, as also becomes clear from the frequent occurrence of reported coverage levels >100% and hence difficult to interpret [33]. Low coverage is problematic, particularly if it is sustained over multiple treatment rounds. We could not assess the importance of this phenomenon in our data, as it appeared difficult to match coverage data from subsequent years at community level and to match them to the mf prevalence data.

The high variation in mf prevalence after a given number of treatment rounds within districts, as observed in Ghana (this study) and elsewhere [5-7] complicates decision making. If communities with high residual mf prevalence are by chance not included in surveys, MDA may be stopped prematurely with danger of resurgence [5]. This could be prevented by targeting pre-TAS surveys to communities at high risk of residual transmission. High risk may occur due to programmatic or demographic factors [12], including migration during treatment period, treatment fatigue, high numbers of middle aged women (child bearing age; majority not taking drug due to pregnancy) etc. Other local factors contributing to transmission may be high biting rate of the mosquitoes and behavior of residents that influence exposure to mosquito bites [32,34]. Moreover, TAS is designed to cover a larger geographical area, with the hope that pockets with residual transmission would be identified in TAS surveys. However, it is unclear whether TAS is sufficiently sensitive to pick up such pockets since not all communities within the evaluation unit (district) are sampled, furthermore, some have reported transmission ongoing in spite of passing TAS [5,6]. Thus, the validity of TAS for longer-term post-MDA surveillance requires further investigation [35].

We could not assess this in the current study, as most districts are usually not resurveyed shortly after stopping MDA. We have data for only 6 communities surveyed after stopping MDA, and all showed mf prevalence <1%.

Our data had some limitations. Firstly, we only considered mf data, as antigenaemia prevalence data were not always collected. Survey sites (apart from spot-check sites) were not randomly chosen, but rather based on previous results and location, and most mf sites have been surveyed only once. The low number of persons sampled combined with less sensitive mf tests in early years makes the mf prevalence observed in early years less reliable (wide 95% CI, data for baseline shown in supplementary data, Table S1). Also, the selection of participants for night blood collection in each community was also not random since some households were more likely to attend than others. This is particularly problematic if those not participating in surveys are also more likely not to participate in MDA, resulting in biased and possibly flattered mf prevalence estimates.

## Conclusions and recommendations

The Ghana Filariasis Elimination programme has had large impact, reducing mf prevalence <1% in 81/98 endemic districts. The remaining 17 districts still need MDA but also seem to be approaching elimination. There was variation in the required treatment rounds between and within districts. Stopping MDA must be done with caution, taking into account the risk that communities with residual transmission remain which could present a source for the resurgence of infection after stopping MDA. Monitoring at the community level is required to be maintained to sustain the gains that have already been made towards elimination of LF in Ghana.

## Acknowledgements

We acknowledge the support from the chiefs, opinion leaders and the community drug distributors in the various communities for mobilizing participants during treatment and mf surveys. We also acknowledge the contributions from the NTD Programme team of the Ghana Health Services (GHS) for planning, implementing and coordinating all field activities and collation of data made available for this article.

## Supporting Information

### S1 Checklist: STROBE Checklist

**Table S1:**
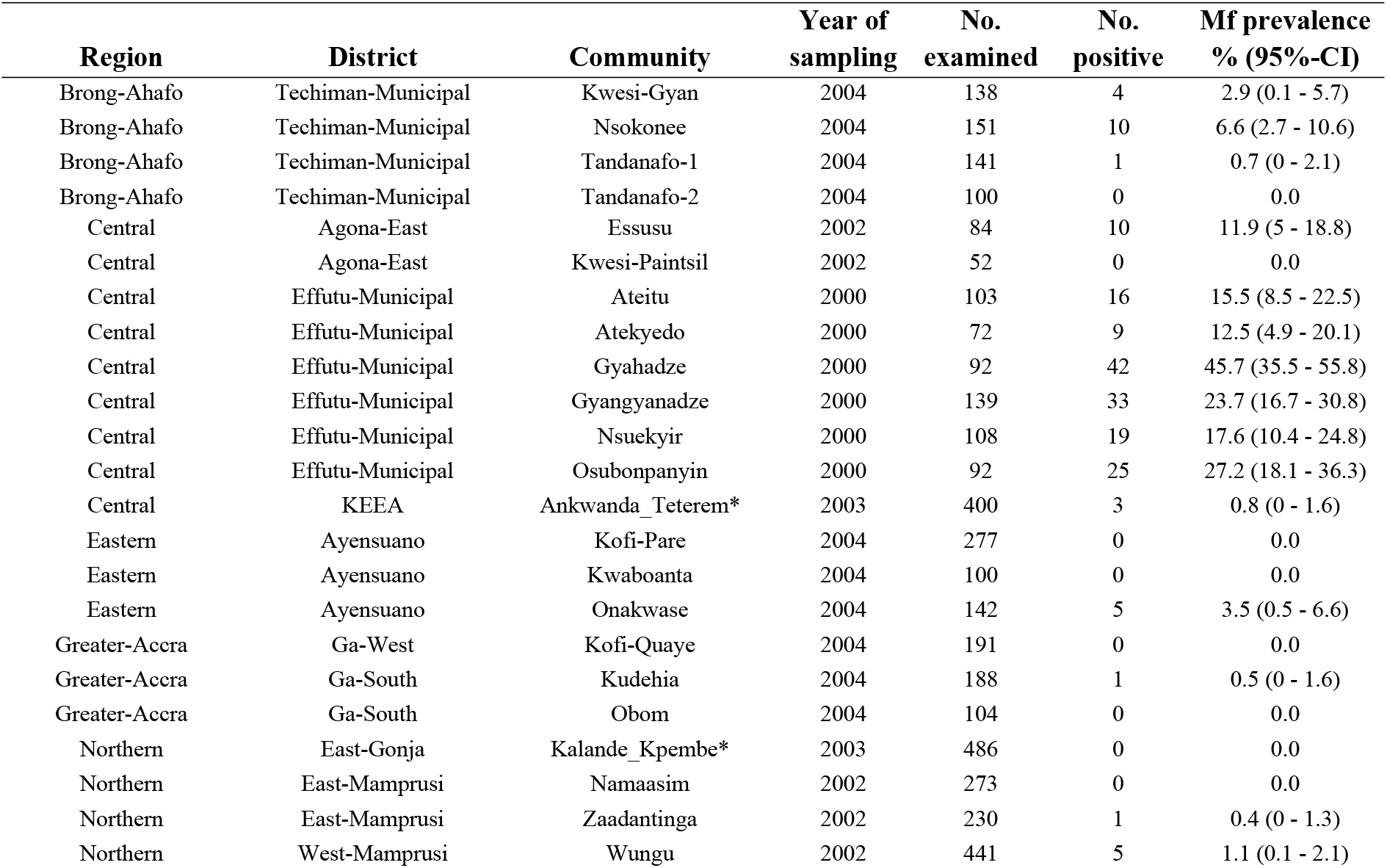
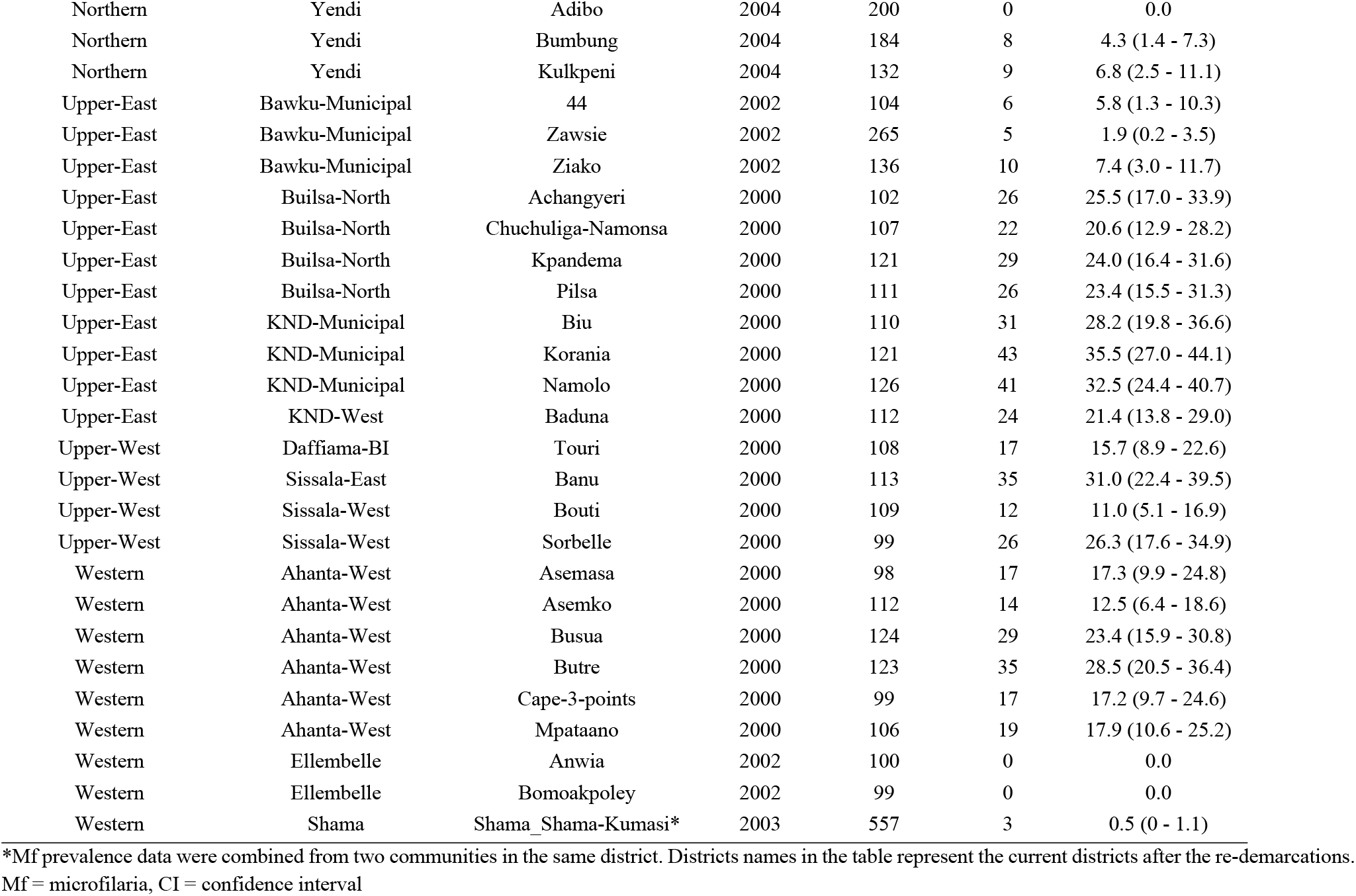
Community-based Mf prevalence at baseline in year 2000 - 2004.

**Table S2:** Overview of MDA implementation and progress towards elimination by district in Ghana, for the 98 districts that were identified as endemic.

| District | District number | Region | District population (2010) | Mf data available - Time post 1st treatment (year) | Baseline mf prevalence (range) | MDA start year* | TAS-1 (Year of last treatment) | Total no. of treatment rounds received by 2016 | TAS-2 (year) | TAS-3 (year) | Hotspot (2016) MDA on-going |
| --- | --- | --- | --- | --- | --- | --- | --- | --- | --- | --- | --- |
| Abura Asebu Kwamankese (AAK) | 28 | Central | 117,185 | 8 | - | 2003 | 2014 | 11 | 2016 | Not yet | - |
| Accra Metro | 96 | Greater Accra | 1,848,614 | 7 | - | 2006 | 2014 | 8 | 2016 | Not yet | - |
| Agona East | 11 | Central | 85,920 | 0,1 | 6 (0 - 11.9) | 2002 | 2010 | 9 | 2012 | 2015 | - |
| Agona West Municipal | 12 | Central | 115,358 | - | - | 2002 | 2010 | 9 | 2012 | 2015 | - |
| Ahanta West | 1 | Western | 106,215 | 0,3,7,12,14 | - | 2000 | Not yet | 16 | Not yet | Not yet | Yes |
| Ajumaku Enyan Essiam (AEE) | 29 | Central | 138,046 | 8 | - | 2003 | 2014 | 11 | 2016 | Not yet | - |
| Akrumfi | 30 | Central | 52,231 | - | - | 2003 | 2014 | 11 | Not yet | Not yet | - |
| Akwapim South | 71 | Eastern | 123,501 | 6 | - | 2005 | 2014 | 9 | 2016 | Not yet | - |
| Aowin | 55 | Western | 138,415 | 7 | 19.5 (12.5 - 28.5) | 2004 | 2014 | 10 | 2016 | Not yet | - |
| Asikuma Odoben Brakwa (AOB) | 56 | Central | 112,706 | 7 | - | 2004 | 2014 | 10 | 2016 | Not yet | - |
| Assin North | 57 | Central | 161,341 | - | - | 2004 | 2014 | 10 | 2016 | Not yet | - |
| Assin South | 58 | Central | 104,244 | 7 | - | 2004 | 2014 | 10 | 2016 | Not yet | - |
| Awutu Senya East Municipal | 2 | Central | 108,422 | - | - | 2001 | 2010 | 10 | 2012 | 2015 | - |
| Awutu Senya West | 3 | Central | 86,884 | 6 | - | 2001 | 2010 | 10 | 2012 | 2015 | - |
| Ayensuano | 72 | Eastern | 77,193 | 0,6 | - | 2005 | 2014 | 9 | 2016 | Not yet | - |
| Bawku Municipal | 31 | Upper East | 217,791 | 0,9 | 5 (1.9 - 5.8) | 2003 | 2014 | 11 | 2016 | Not yet | - |
| Bawku West | 32 | Upper East | 94,034 | 6,10 | - | 2003 | 2014 | 11 | Not yet | Not yet | - |
| Binduri | 33 | Upper East | 61,576 | - | - | 2003 | 2014 | 11 | 2016 | Not yet | - |
| Bole | 68 | Northern | 61,593 | 7,10 | - | 2004 | Not yet | 12 | Not yet | Not yet | Yes |
| Bolgatanga Municipal | 15 | Upper East | 131,550 | 1,2,7,11 | - | 2002 | 2015 | 13 | Not yet | Not yet | - |
| Bongo | 16 | Upper East | 84,545 | 1,2,7,11 | - | 2002 | 2015 | 13 | Not yet | Not yet | - |
| Builsa North | 7 | Upper East | 56,477 | 0,2,6,11 | 23.4 (20.6 - 25.5) | 2001 | 2015 | 14 | Not yet | Not yet | - |
| Builsa South | 8 | Upper East | 36,514 | 6,12 | - | 2001 | 2015 | 14 | Not yet | Not yet | - |
| Bunkprugu Yunyoo | 73 | Northern | 122,591 | 7 | - | 2005 | 2014 | 9 | 2016 | Not yet | - |
| Cape Coast | 59 | Central | 169,894 | 7 | - | 2004 | 2014 | 10 | 2016 | Not yet | - |
| Central Gonja | 74 | Northern | 87,877 | 6 | - | 2005 | 2014 | 9 | 2016 | Not yet | - |
| Chereponi | 34 | Northern | 53,394 | 8 | - | 2003 | 2014 | 11 | 2016 | Not yet | - |
| Daffiama Busie Issa | 17 | Upper West | 32,827 | 0,1,2,7,11 | - | 2002 | 2015 | 13 | Not yet | Not yet | - |
| East Gonja | 60 | Northern | 135,450 | 7 | 0 | 2004 | 2014 | 10 | 2016 | Not yet | - |
| East Mamprusi | 35 | Northern | 121,009 | 0,9 | 0.22 (0 - 0.4) | 2003 | 2014 | 11 | 2016 | Not yet | - |
| Effutu Municipal | 4 | Central | 68,597 | 0,2,6,13 | 23.7 (12.5-45.7) | 2001 | 2010 | 10 | 2012 | 2015 | - |
| Ellembelle | 20 | Western | 87,501 | 0,7,10 | - | 2002 | Not yet | 14 | Not yet | Not yet | Yes |
| Ga Central | 75 | Greater Accra | 117,220 | - | - | 2005 | 2014 | 9 | 2016 | Not yet | - |
| Ga East | 76 | Greater Accra | 259,668 | 6 | - | 2005 | 2014 | 9 | 2016 | Not yet | - |
| Ga South | 77 | Greater Accra | 485,643 | 0,6 | 0.27 (0 - 0.53) | 2005 | 2014 | 9 | 2016 | Not yet | - |
| Ga West | 78 | Greater Accra | 262,742 | 0,6 | 0 | 2005 | 2014 | 9 | 2016 | Not yet | - |
| Garu Tempane | 79 | Upper East | 130,003 | 7 | - | 2005 | 2014 | 9 | 2016 | Not yet | - |
| Gomoa East | 13 | Central | 207,071 | 9 | - | 2002 | 2014 | 12 | 2016 | Not yet | - |
| Gomoa West | 14 | Central | 135,189 | - | - | 2002 | 2014 | 12 | 2016 | Not yet | - |
| Gushiegu | 36 | Northern | 111,259 | - | - | 2003 | 2014 | 11 | 2016 | Not yet | - |
| Jirapa | 21 | Upper West | 88,402 | 1,2,7,11 | - | 2002 | Not yet | 14 | Not yet | Not yet | Yes |
| Jomoro | 37 | Western | 150,107 | 8 | - | 2003 | 2014 | 11 | 2016 | Not yet | - |
| Karaga | 38 | Northern | 77,706 | 8 | - | 2003 | 2014 | 11 | Not yet | Not yet | - |
| KEEA | 61 | Central | 144,705 | 7 | 0.75 | 2004 | 2014 | 10 | 2016 | Not yet | - |
| KND-Municipal | 9 | Upper East | 109,944 | 0,2,6,11,13 | 32.1 (28.1 - 35.5) | 2001 | 2015 | 14 | Not yet | Not yet | - |
| KND-West | 10 | Upper East | 70,667 | 0,2,4,6,11,13 | 21.4 | 2001 | Not yet | 15 | Not yet | Not yet | Yes |
| Kpandai | 62 | Northern | 108,816 | - | - | 2004 | 2014 | 10 | 2016 | Not yet | - |
| Kumbungu | 39 | Northern | 39,341 | 6 | - | 2003 | 2014 | 11 | 2016 | Not yet | - |
| La Dade Kotopon | 97 | Greater Accra | 183,528 | 7 | - | 2006 | 2014 | 8 | 2016 | Not yet | - |
| La Nkwantanang Madina | 80 | Greater Accra | 111,926 | - | - | 2005 | 2014 | 9 | 2016 | Not yet | - |
| Lambussie Karni | 22 | Upper West | 51,654 | 1,2,7,12 | - | 2002 | Not yet | 14 | Not yet | Not yet | Yes |
| Lawra | 23 | Upper West | 100,929 | 1,2,7,11 | - | 2002 | Not yet | 14 | Not yet | Not yet | Yes |
| Ledzokuku krowor | 98 | Greater Accra | 227,932 | 7 | - | 2006 | 2014 | 8 | 2016 | Not yet | - |
| Mamprugu Moaduri | 40 | Northern | 46,894 | 6 | - | 2003 | 2014 | 11 | 2016 | Not yet | - |
| Mfantseman | 41 | Central | 196,563 | 8 | - | 2003 | 2014 | 11 | 2016 | Not yet | - |
| Mion | 81 | Northern | 81,812 | - | - | 2005 | 2014 | 9 | 2016 | Not yet | - |
| Mpohor | 42 | Western | 123,996 | - | - | 2003 | 2014 | 11 | 2016 | Not yet | - |
| Nabdam | 92 | Upper East | 33,826 | 8 | - | 2005 | Not yet | 11 | Not yet | Not yet | Yes |
| Nadowli | 18 | Upper West | 94,388 | 7,11 | 15.7 | 2002 | 2015 | 13 | Not yet | Not yet | - |
| Nandom | 24 | Upper West | 46,040 | - | - | 2002 | Not yet | 14 | Not yet | Not yet | Yes |
| Nanumba North | 43 | Northern | 141,584 | 6 | - | 2003 | 2014 | 11 | 2016 | Not yet | - |
| Nanumba South | 44 | Northern | 93,464 | - | - | 2003 | 2014 | 11 | 2016 | Not yet | - |
| North Gonja | 69 | Northern | 43,547 | 10 | - | 2004 | Not yet | 12 | Not yet | Not yet | Yes |
| Nsawam Adoagyiri | 82 | Eastern | 86,000 | - | - | 2005 | 2014 | 9 | 2016 | Not yet | - |
| Nzema East | 25 | Western | 60,828 | 7,10,12 | 0 | 2002 | Not yet | 14 | Not yet | Not yet | Yes |
| Prestea Huni Valley | 45 | Western | 159,304 | - | - | 2003 | 2014 | 11 | 2016 | Not yet | - |
| Pusiga | 46 | Upper East | 57,677 | - | - | 2003 | 2014 | 11 | 2016 | Not yet | - |
| Saboba | 47 | Northern | 65,706 | 8 | - | 2003 | 2014 | 11 | 2016 | Not yet | - |
| Sagnerigu | 83 | Northern | 148,099 | - | - | 2005 | 2014 | 9 | 2016 | Not yet | - |
| Savelugu Nanton | 48 | Northern | 139,283 | 6 | - | 2003 | 2014 | 11 | 2016 | Not yet | - |
| Sawla Tuna Kalba | 93 | Northern | 99,863 | 6,9 | - | 2005 | Not yet | 11 | Not yet | Not yet | Yes |
| Sekondi Takoradi metro | 63 | Western | 559,548 | 8 | - | 2004 | 2014 | 10 | 2016 | Not yet | - |
| Shama | 64 | Western | 81,966 | 7 | - | 2004 | 2014 | 10 | 2016 | Not yet | - |
| Sissala East | 5 | Upper West | 56,528 | 0,6,11 | 0.54 | 2001 | 2014 | 13 | 2016 | Not yet | - |
| Sissala West | 6 | Upper West | 49,573 | 0,2,6,11 | 31 | 2001 | 2014 | 13 | 2016 | Not yet | - |
| Suaman | 65 | Western | 20,529 | - | 18.6 (11 - 26) | 2004 | 2014 | 10 | 2016 | Not yet | - |
| Suhum | 84 | Eastern | 167,551 | 6 | - | 2005 | 2014 | 9 | 2016 | Not yet | - |
| Sunyani Municipal | 94 | Brong Ahafo | 123,224 | 7,9 | 1.2 (0 - 3.5) | 2005 | Not yet | 11 | Not yet | Not yet | Yes |
| Sunyani West | 95 | Brong Ahafo | 85,272 | 7,9 | - | 2005 | Not yet | 11 | Not yet | Not yet | Yes |
| Talensi | 89 | Upper East | 115,020 | 8 | - | 2005 | 2015 | 10 | Not yet | Not yet | - |
| Tamale Metro | 85 | Northern | 371,351 | 6 | - | 2005 | 2014 | 9 | 2016 | Not yet | - |
| Tarkwa Nsuaem | 49 | Western | 90,477 | 5 | - | 2003 | 2014 | 11 | 2016 | Not yet | - |
| Tatale Sanguli | 50 | Northern | 60,039 | 6 | - | 2003 | 2014 | 11 | 2016 | Not yet | - |
| Techiman Municipal | 90 | Brong Ahafo | 206,856 | 0,7,9 | - | 2005 | 2016 | 11 | Not yet | Not yet | - |
| Techiman North | 91 | Brong Ahafo | 59,068 | 9 | 2.6 (0 - 6.6) | 2005 | 2016 | 11 | Not yet | Not yet | - |
| Tolon | 51 | Northern | 112,331 | 6 | - | 2003 | 2014 | 11 | 2016 | Not yet | - |
| Twifo Ati Mokwa | 66 | Central | 61,743 | - | - | 2004 | 2014 | 10 | 2016 | Not yet | - |
| Twifo Heman Lower Denkyira | 67 | Central | 116,874 | 7 | - | 2004 | 2014 | 10 | 2016 | Not yet | - |
| Upper West Akim | 86 | Eastern | 87,051 | 6 | - | 2005 | 2014 | 9 | 2016 | Not yet | - |
| Wa East | 26 | Upper West | 72,074 | 11 | - | 2002 | Not yet | 14 | Not yet | Not yet | Yes |
| Wa Municipal | 19 | Upper West | 107,214 | 11 | - | 2002 | 2015 | 13 | Not yet | Not yet | - |
| Wa West | 27 | Upper West | 81,348 | 1,2,7,11 | - | 2002 | Not yet | 14 | Not yet | Not yet | Yes |
| Wassa East | 52 | Western | 81,073 | 8 | - | 2003 | 2014 | 11 | 2016 | Not yet | - |
| West Akim Municipal | 87 | Eastern | 195,349 | 6 | - | 2005 | 2014 | 9 | 2016 | Not yet | - |
| West Gonja | 70 | Northern | 84,727 | 7,10 | - | 2004 | Not yet | 12 | Not yet | Not yet | Yes |
| West Mamprusi | 53 | Northern | 168,011 | 0,6 | 1.1 | 2003 | 2014 | 11 | 2016 | Not yet | - |
| Yendi | 88 | Northern | 199,592 | 0,6 | 3.7 (0 - 6.8) | 2005 | 2014 | 9 | 2016 | Not yet | - |
| Zabzugu | 54 | Northern | 123,854 | 6 | - | 2003 | 2014 | 11 | 2016 | Not yet | - |

\* All communities in each district were expected to be treated in the same year MDA started, so the geographical coverage at district level is 100% right from the start. So far, all TAS surveys were passed (no failures). The year in which TAS surveys were done, were the same year of last MDA. Districts names in the table represent the current districts after the re-demarcations. Need to explain all abbreviations here

**Figure S1:**
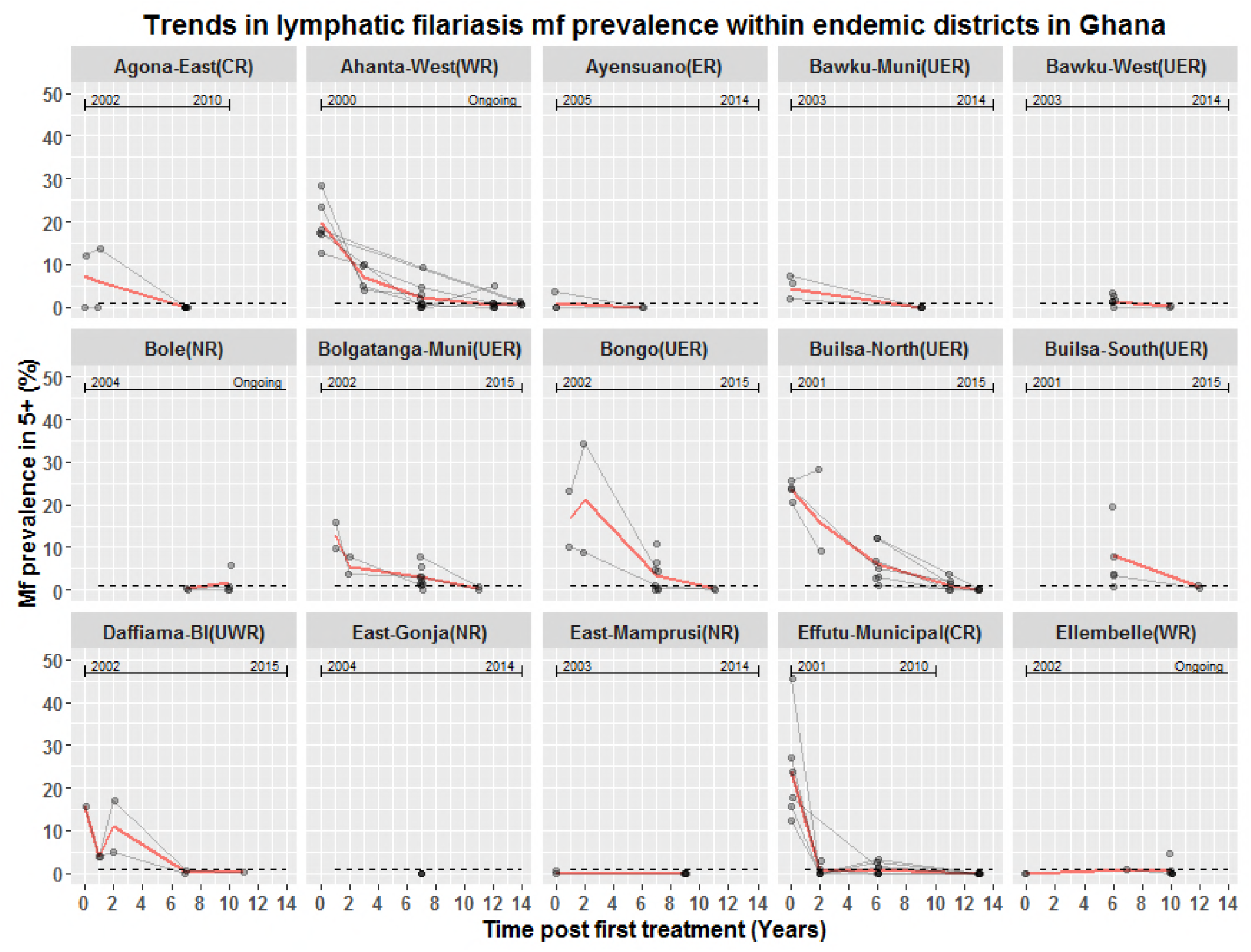
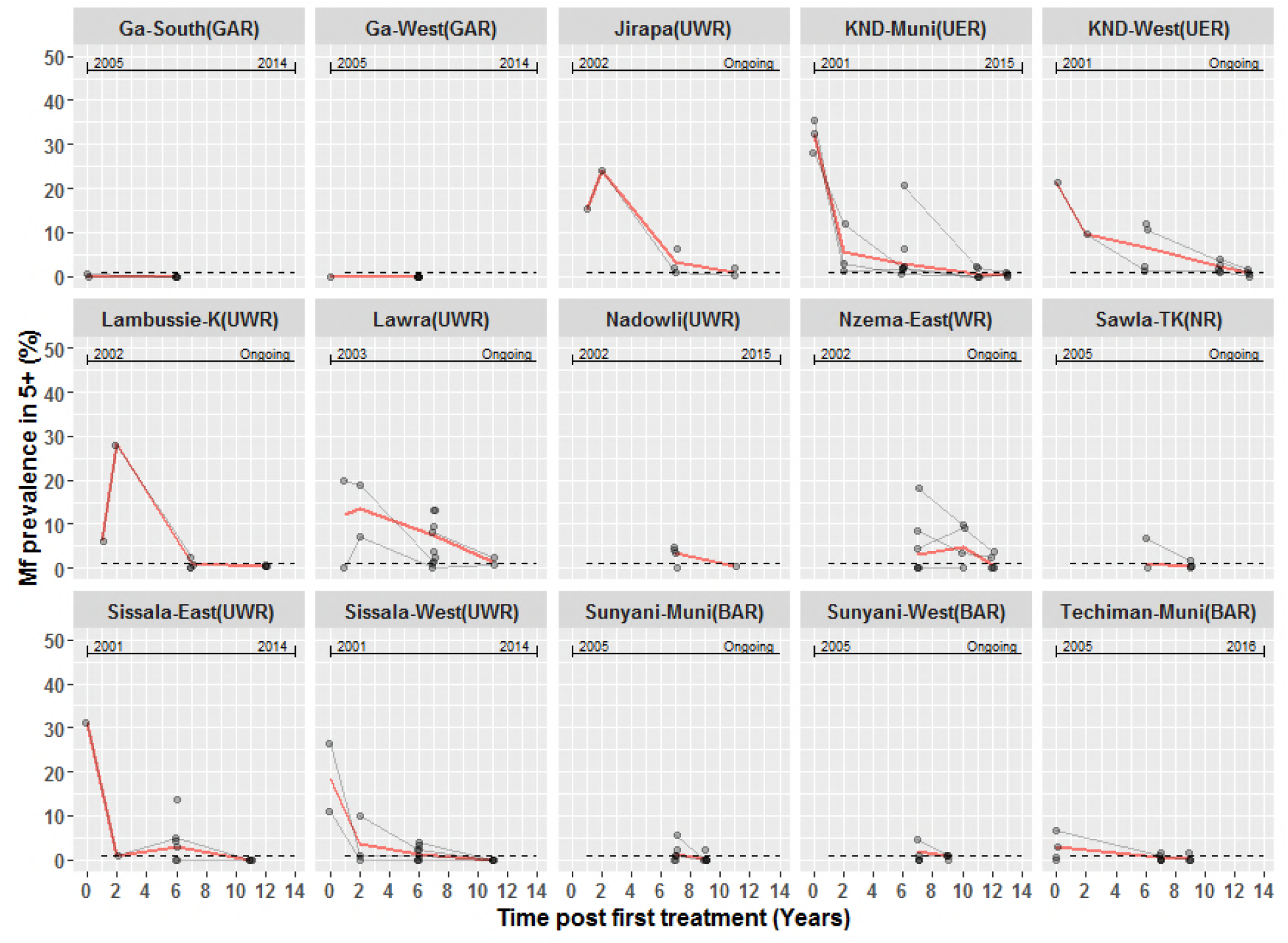
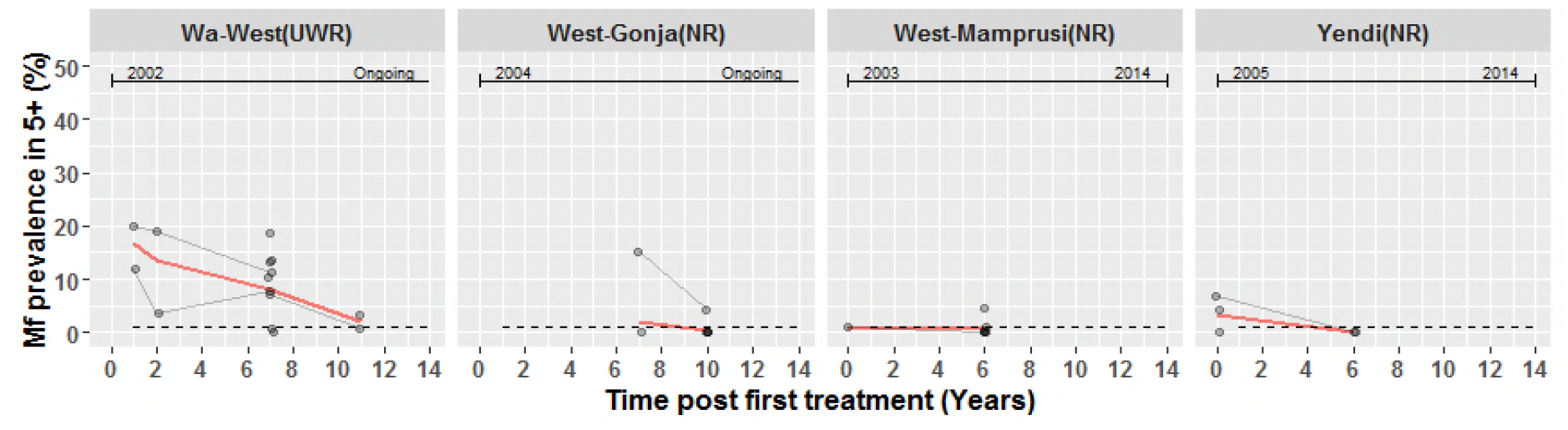
Mf prevalence distribution overtime in 34/98 districts with data for multiple time points. Each panel represents a district which was sampled multiple times during the survey. The red continuous line represent the average prevalence in the district at each time point estimated from communities sampled within the same district. The bullets are community mf prevalence at each time point and multiple observations from the same community are connected through thin grey lines. The dashed horizontal lines represent the threshold mf prevalence of 1%. The line at the upper part of each panel with the text indicate the year MDA started and when it ended or if ongoing by September 2016. The suffix attached to district names represent the region in which district is located as follows: Brong Ahafo (BAR), Central (CR), Eastern (ER), Greater Accra (GAR), Northern (NR), Upper East (UER), Upper West (UWR) and Western (WR). Districts names in the figure represent the current districts after the re-demarcations.

**Figure S2:**
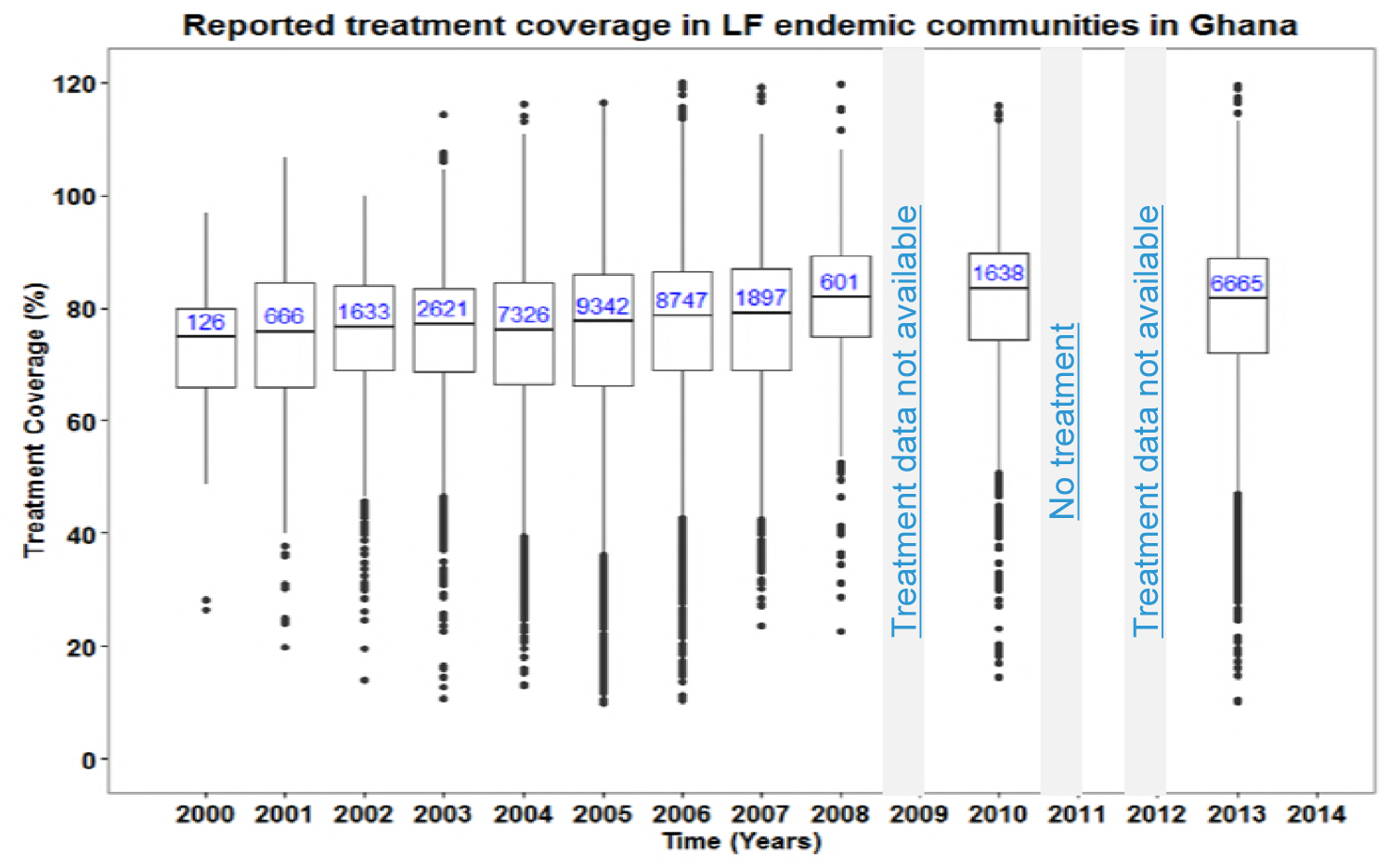
Reported treatment coverage in treated communities in Ghana. The box at each time point represents the interquartile range of coverage and the thick horizontal lines across each box represent the median coverage. The bullets outside each box (above or below) represent the outliers and are calculated as 1.5 times the interquartile range above or below the ends of the box (25^th^ and 75^th^ percentile). The vertical lines (whiskers) extend to the first value (coverage) before the outlier cut-off and where there are no outliers, they represent the minimum and maximum coverage at each time point. The numbers in the boxes are the total number of communities treated at each time point. There was no treatment offered in 2011 due to some challenges; 2009 and 2012 treatment data not available.

We acknowledge the fact that these reported coverage may not be very reliable [33] since some areas reported coverage more than 100% (Figure S2). Our data showed a relatively high and constant coverage over time. Some of the flaws may probably be caused by community registers used in estimating the population not being updated, resulting in high coverage and more individuals treated than the existing population in some cases. Treatment coverage may also be overestimated because community directed distributors (CDD) are recommended to treat by direct observed treatment (DOT) but they may not have observed all persons but may record as treated. On the other hand the incidental low coverage could also be due to under-reporting especially where individuals treated by the CDD (who sometimes kept leftover drugs from previous treatments) before or after the GHS treatment period when treatment books were in the possession of the GHS sub-district office.

